# Role of Inferior Frontal Junction (IFJ) in the Control of Feature vs Spatial Attention

**DOI:** 10.1101/2020.11.04.368993

**Authors:** Sreenivasan Meyyappan, Abhijit Rajan, George R Mangun, Mingzhou Ding

**Affiliations:** J. Crayton Pruitt Family Department of Biomedical Engineering, University of Florida, Gainesville, FL 32611; Center for Mind and Brain, University of California, Davis, CA 95618; Departments of Psychology and Neurology, University of California, Davis, CA 95618

## Abstract

Feature-based attention refers to preferential selection and processing of items and objects based on their non-spatial attributes such as color or shape. While it is intuitively an easier form of attention to relate to in our day to day lives, the neural mechanisms of feature-based attention are not well understood. Studies have long implicated the dorsal attention network as a key control system for voluntary spatial, feature and object-based attention. Recent studies have expanded on this model by focusing on the inferior frontal junction (IFJ), a region in the pre-frontal cortex to be the source of feature attention control, but not spatial attention control. However, the extent to which IFJ contributes to spatial attention remains a topic of debate. We investigated the role of IFJ in the control of feature versus spatial attention in a cued visual spatial (attend left or right) and feature attention (attend red or green) task using fMRI. Analyzing single-trial cue-evoked fMRI responses using univariate GLM and multi-voxel pattern analysis (MVPA), we observed the following. First, the univariate BOLD activation responses yielded no significant differences between feature and spatial cues. Second, MVPA analysis showed above chance level decoding in classifying feature attention (attend-red vs. attend-green) in both the left and right IFJ, whereas during spatial attention (attend-left vs. attend-right) decoding was at chance. Third, while the cue-evoked decoding accuracy was significant for both left and right IFJ during feature attention, target stimulus-evoked neural responses were not different. Importantly, only the connectivity patterns from the right IFJ was predictive of target-evoked activity in visual cortex (V4); this was true for both left and right V4. Finally, the strength of this connectivity between right IFJ and V4 (bilaterally) was found to be predictive of behavioral performance. These results support a model where the right IFJ plays a crucial role in top down control of feature but not spatial attention.

## INTRODUCTION

Attention can be captured by salient external stimuli (exogenous attention) or deployed voluntarily according to behavioral goals (endogenous attention). Endogenous attention to sensory events can be deployed based on their relevant stimulus attributes such as locations, features (e.g., color) or forms (e.g. objects), as well their conjunctions. Attentional cuing paradigms (e.g., Posner, 1981) are especially effective for investigating the neural mechanisms of attention control to these stimulus attributes and can reveal information about both attentional control processes and their effect on stimulus processing.

In typical cueing paradigms, each trial starts with a cue that defines the behaviorally relevant attribute of a stimulus (target) that will appear after a delay of milliseconds or seconds. The cue enables the subject to engage preparatory attention for the relevant stimulus attribute, such as its location (e.g., Harter et al., 1989; Mangun and Hillyard, 1991; Corbetta et al., 2000; Hopfinger et al., 2000), color, motion or other features (Giesbrecht et al., 2003; Snyder and Fox, 2010), or higher-level object properties (e.g., Noah et al., 2020). When the anticipated target stimulus appears, the subject is required to execute a task and make a response according to predefined rules (behavioral goals).

Two decades of research using fMRI have shown that in the time period following the cue but in advance of the target stimulus (the cue-to-target interval), the dorsal attention network (DAN), principally comprising the intraparietal sulcus (IPS) and frontal eye fields (FEF), is activated irrespective of whether attention is directed to a spatial location or a non-spatial stimulus feature (Liu et al., 2003; Giesbrecht et al., 2003; Slagter et al., 2007; Bichot et al., 2015). This has led to the notion that the DAN is an important neural network supporting the top-down control of attention regardless of the to-be-attended stimulus attribute. More specifically, in the cue-to-target interval, the DAN is thought to maintain the attentional set and issue top-down control signals to sensory-specific cortex in order to bias sensory processing so that the attended information is facilitated and irrelevant stimulus inputs are suppressed (e.g., Corbetta et al., 2008; Wang et al., 2016).

Regions outside the classically defined DAN have also been implicated in attentional control. The inferior frontal junction (IFJ) is such an area. IFJ is a prefrontal structure situated at the confluence of the precentral, superior, and inferior frontal sulcus (Derrfuss et al., 2009; Zanto et al., 2010; Baldauf and Desimone, 2014; Zhang et al., 2018). Earlier work on IFJ function focused on its role in cognitive control, using paradigms such as task switching (Brass and von Cramon 2004; Brass et al., 2005; Derfuss et al., 2005) and Stroop (Neumann et al., 2005; Derfuss et al., 2005). A more recent line of inquiry has focused on IFJ’s role in attention control (Nobre et al., 1997; Zanto et al., 2010 Asplund et al., 2010, Baldauf and Desimone 2014; Bichot et al., 2015; Bichot et al., 2019). It was shown that IFJ, along with the nodes in the DAN, is activated by spatial attention cues (Giesbrecht et al., 2003; Asplund et al., 2010), leading to the notion that IFJ is involved in the control of spatial attention and that it may even be a part of the DAN (Corbetta et al., 2008). In contrast, however, using an experimental design with long cue-to-target intervals, Tamber-Rosenau et al. (2018; see also Asplund et al., 2010) challenged this notion, showing that during spatial attention the IFJ activity following a cue was short-lived, suggesting that it was more involved in interpreting the cues and transitioning attentional orienting, rather than controlling sustained attention to spatial locations *per se*.

Besides spatial attention control, increasingly, IFJ’s role in the control of non-spatial feature attention is being investigated (Zanto et al., 2010; Baldauf and Desimone 2014; Bichot et al., 2015; Zhang et al., 2018; Bichot et al., 2019; Gong and Liu 2020). Zanto et al. (2010), using fMRI and functional connectivity, showed that IFJ selectively modulated activity within visual sensory areas of V4 and MT when subjects attended to color and motion, respectively. Baldauf and Desimone (2014), using MEG and fMRI, identified IFJ as a key modulator of activity in the FFA when subjects attended to faces, and the PPA when subjects attended to houses while viewing a stream of face-house compound stimuli in a one-back task. They reported that the IFJ-sensory cortex coupling, indexed by gamma frequency synchrony, was sustained in the time period between stimuli. In both Zanto et al. (2010) and Baldauf and Desimone (2014), the subjects attended to different feature domains (e.g., motion vs. color; houses vs. faces), but not to different stimuli within a given feature domain. Thus, to what extent IFJ activity codes different attended information within each feature domain (e.g., male vs. female faces) remains to be elucidated. Gong and Liu (2020) addressed this question by investigating the neural representations in IFJ when attention is directed to different colors or motion directions and finding decodable neural activity differences in IFJ within each attended feature domain using MVPA. Their paradigm, however, differed from the cueing paradigm mentioned in the foregoing, in that the stimulus was present throughout the trial, and the subject detected changes in the stimulus. Thus, attention control and attention selection of stimulus are not well separated. Shedding light on IFJ’s role in controlling feature attention in the absence of visual stimulation remains a goal to be attained.

The hemispheric differences in IFJ’s control of attention are also a debated issue. Zanto et al. (2010) showed that while attending to motion elicits increase in bilateral IFJ connectivity with MT cortex, attending to color evokes only increased right IFJ (rIFJ) connectivity with V4. Baldauf et al. (2014) extended the findings of Zanto et al. (2010) to the domain of object attention by showing that the functional connectivity between right IFJ and PPA was increased when subjects attended houses in compound face-house stimuli whereas the functional connectivity between right IFJ and FFA was increased when subjects attended faces. Lee and Geng (2017) used a paradigm in which a morphed face (between two famous actors’ faces) was presented, and the subjects were required to categorize the stimulus as one of the two actors faces. Using functional MRI and representational similarity analyses, they revealed the behavioral categorization of the morphed faces was correlated with neural activity in lateral prefrontal cortex and the ventral temporal cortex (e.g., fusiform face area), but only the right lateral ventral prefrontal cortex coded individual differences in categorization behavior. A more recent study by Zhang et al. (2018) presented lateralized stimuli (left visual field vs right visual field) to subjects and applied effective connectivity to fMRI. They found higher contralateral, relative to ipsilateral, IFJ-visual cortex connectivity but did not report on the difference between left and right IFJ. In contrast, however, Gong and Liu (2020), described above, reported no specific hemispheric differences within IFJ in terms of decoding accuracies. Thus, while there is evidence for right hemisphere specialization for the role of IFJ in attentional processing, there also remains uncertainty, requiring further investigation.

Multivariate pattern analysis (MVPA) techniques are known to be more sensitive at distinguishing nuanced pattern variations driven by subtle differences in experimental conditions (Haxby et al., 2001; Kamitani and Tong 2005; Liu and Hou 2013; Gong and Liu 2020). As noted, the connectivity analyses are also illuminating, and therefore combining functional connectivity and MVPA decoding approaches in the same cuing experiment could shed further light on the role of the IFJ in the control of attention.

In the present study, fMRI data was recorded from human participants performing a cued spatial/feature attention experiment. At the beginning of each trial an auditory cue instructed the participant to covertly pay attention to either one of two spatial locations (left versus right visual field), referred to as spatial trials, or one of two colors (red versus green), referred to as feature or color trials. Following a random delay, two rectangular stimuli appeared, one in each visual field, and the subjects reported the orientation of the rectangle either in the attended location (spatial trials) or having the attended color (feature trials). We note that, by including feature cues and spatial cues in the same experiment, we were able to better compare the role of IFJ in the control of these two types of attention. Analytically, MVPA methods were applied along with conventional univariate analysis, to examine more comprehensively the presence and specificity of attention control signals in IFJ during anticipatory feature vs spatial attention. An additional issue explored is whether IFJ is mainly engaged in the control of attention (cue evoked activity) or whether it is also engaged in attention selection (target evoked activity).

## METHODS

### Overview

Twenty (5 female) right-handed graduate and undergraduate students took part in the study. The participants reported no prior history of neuro-psychological disorders and provided written informed consent before taking part in the study. The experimental protocol was approved by the Institutional Review Board (IRB) at the University of Florida. Data from this experiment has been used to investigate different questions in our previously published work (Rajan et al., 2019; Rajan et al., under review).

### Procedure

The time-course of a typical trial is illustrated in Figure 1. Participants were expected to fixate at the central cross. Two sets of dots indicated the two peripheral locations, 3.6 degrees lateral to the upper left and upper right of the fixation cross. Start of the trial was signaled by an auditory cue, which directed the participants to covertly direct their attention to either a spatial location (“left” or “right”) or to a color (“red” or “green”). Following a delay period, varied randomly from 3000 ms to 6600 ms, two colored rectangles (red or green) were presented for a duration of 200 ms, with one in each of the two peripheral locations. The subject’s task was to report the orientation of the rectangle (target) appearing at the cued location or having the cued color, and to ignore the other rectangle (distractor). For feature or color trials, the two rectangles displayed were always of the opposite color; for spatial trials, the two rectangles were either of the same color or of the opposite color. On 8% of the trials (invalid trials), only one rectangle was displayed, which was either not in the cued location for spatial trials or not having the cued color for color trials, and the participants were required to report the orientation of that rectangle. These invalidly cued trials were included to measure the behavioral benefits of attentional cuing (Posner 1980). An inter-trial interval, which was varied randomly from 8000 ms to 12800 ms following the target onset, elapsed before the start of the next trial.

**Figure 1.**
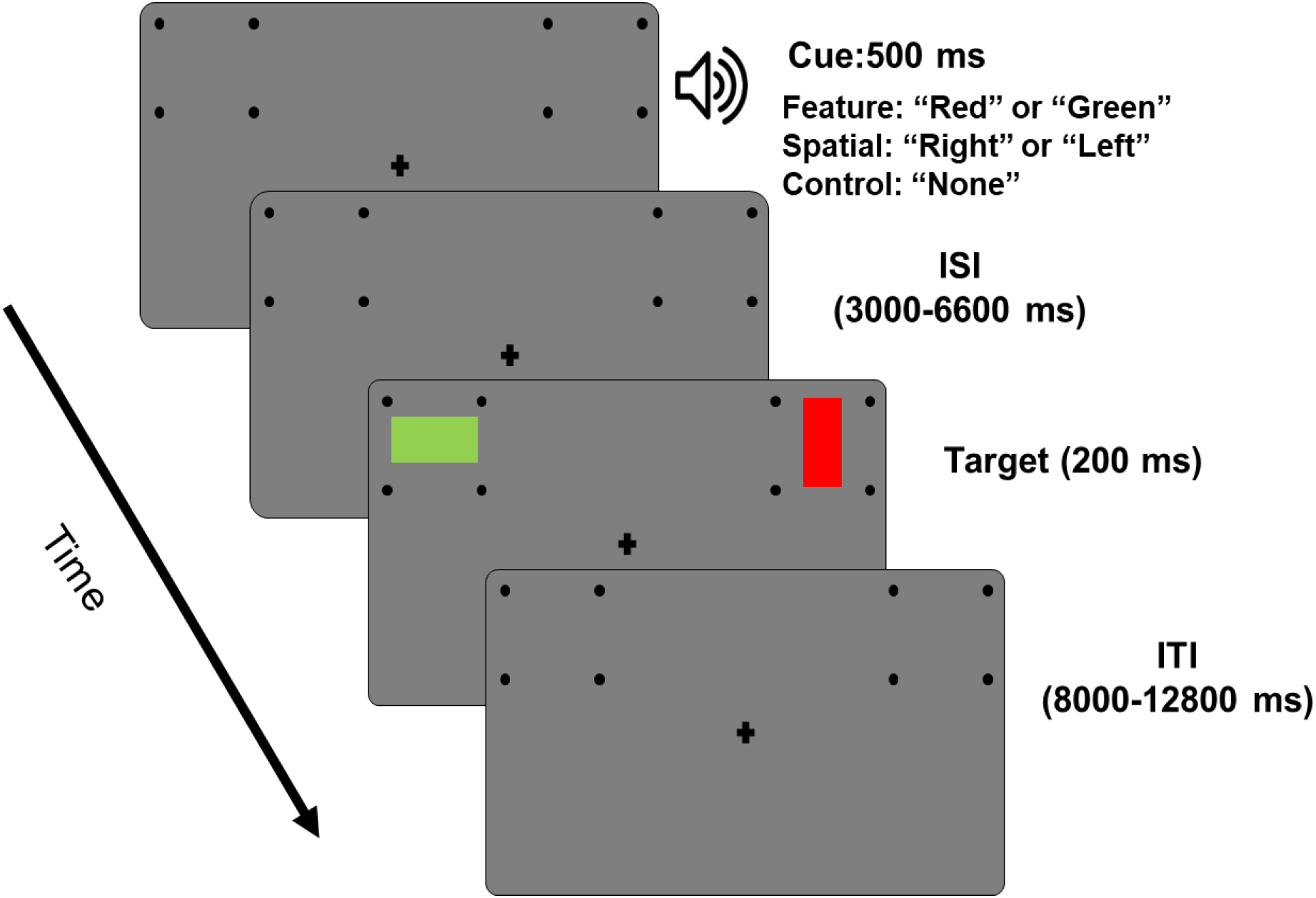
Experimental paradigm. Each trial starts with an auditory cue (500ms) instructing the subject to covertly attend to a spatial location (“left” or “right”) or to a color (“red” or “green”). Following a variable cue-to-target delay (3000-6600ms), two colored rectangles were displayed (100ms), one in each of the two-peripheral location. Participants were asked to report the orientation of the rectangle (horizontal or vertical) displayed in the cued location or having the cued color. On some of the trials (8%) the cues were not valid, i.e., only one target appeared, which was either not at attended location or not having the attended feature, and participants were required to report the orientation of the rectangle. When the none cue was presented (neutral trials), the subject did not prepare attention for spatial or color trials, and instead prepared to discriminate the orientation of the rectangle that was presented over a grey patch. An inter-trial interval, varied randomly from 8000-12800ms following the target onset, elapsed before the start of the next trial. On a portion of trials (20%), the cue was non followed by targets (cue-only trials). Note: MVPA analyses within the DAN for spatial and feature attentional control are reported separately in Rajan et al., (under review).

Trials were organized into blocks, with each block consisting of 25 trials and lasting approximately seven minutes. Each participant completed 10 to 14 blocks over two days. In addition to spatial and color cues, there was a third type of “neutral” cue (the word “none”), which comprised of 20% of the total number of trials and provided no information as to what to attend, but instead warned the subject to prepare to respond to a rectangle’s orientation based on it being placed on a grey patch; these were included to provide behavioral measures comparing focused (spatial or feature) versus neutral attention, but were not analyzed here for BOLD activity. Space and color cues were followed (with delay of 3000-6600 ms) 80% of the time by the two colored rectangles (red or green), while on the remaining 20% of trials the cue appeared but no target followed (cue-only trials).

All the participants went through a training session prior to scanning in which they were introduced to the task and became comfortable performing it. Since the study required participants to maintain central fixation for long durations and pay covert attention to the periphery while fixating, participants were screened based on their ability to maintain eye fixation at the center of the monitor throughout the training session. An SR Research EyeLink 1000 eye tracker system was used for that purpose. At the end of training session behavioral accuracy above 70% was considered satisfactory to enroll them in the fMRI study.

### Functional MRI acquisition and preprocessing

A 3T Philips Achieva scanner with a 32-channel head coil (Philips Medical Systems, the Netherlands) was used to collect functional magnetic resonance imaging (fMRI) data. The echo-planar imaging (EPI) sequence parameters were: repetition time (TR), 1.98 s; echo time, 30 ms; flip angle, 80°; field of view, 224 mm; slice number, 36; voxel size, 3.5 × 3.5 × 3.5 mm; matrix size, 64 × 64. The slices were oriented parallel to the plane connecting the anterior and posterior commissures.

The fMRI BOLD (Blood Oxygen Level Dependent) data were processed using statistical parametric mapping toolbox (SPM-12) using custom scripts written in MATLAB. Preprocessing steps included slice timing correction, realignment, spatial normalization, and smoothing. Slice timing correction was carried out using sinc interpolation to correct for differences in slice acquisition time within an EPI volume. The images were then spatially realigned to the first image of each session by a 6-parameter rigid body spatial transformation to account for head movement during acquisition. Each participant’s images were then normalized and registered to the Montreal Neurological Institute (MNI) space. All images were further resampled to a voxel size of 3 × 3 × 3 mm, and spatially smoothed using a Gaussian kernel with 7 mm full width at half maximum. Slow temporal drifts in baseline were removed by applying a high-pass filter with cutoff frequency set at 1/128 Hz.

### Univariate fMRI activation analysis – cue evoked

The general linear model (GLM) method, as implemented in the Statistical Parametric Mapping (SPM) toolbox, was used to analyze the BOLD responses to stimuli. Eight task-related events were included in the GLM analysis as regressors. Five of them were used to model the cue related BOLD activity; only trials with correct responses were included. We used two additional regressors to account for BOLD responses evoked by target stimuli; one for valid and one for invalidly cued targets; in our experimental design, the trials could be either validly cued or invalidly cued, and the subject was expected to respond to both. Finally, one regressor was used to model the trials with incorrect responses.

Group-level cue-evoked fMRI activations were obtained by a parametric one-sample t-test and the threshold for significance was set to p < 0.05 after correcting for multiple comparisons by the false discovery rate (FDR) method.

### Region-of-interest (ROI) definition

Bilateral IFJ ROIs were defined by using the group level cue-evoked BOLD activation. Activation evoked by spatial and feature cues were subjected to a statistical threshold of p<0.05 (FDR correction). Voxels meeting this threshold requirement and lying in the proximity of previously published stereotypical co-ordinates of IFJ (Brass et al., 2005) were taken to be the IFJ ROIs used in this study. The V4 ROI was defined according to a recently published probabilistic atlas of the visual cortex (Wang et al., 2015).

### Estimation of single-trial BOLD response

In addition to the conventional BOLD activity analysis, we also applied multivoxel pattern analysis (MVPA) to the data to examine the differences in patterns evoked by cues and by targets. Patterns of activity evoked by cues and targets, which comprised the main events during a trial, were analysed to address questions in attentional control (cue evoked) and in attentional selection (target evoked). Since MVPA is performed at the single trial level, a beta series regression method (Rissman et al., 2004) was used to estimate BOLD response on each trial in every voxel. In this method, cues, and targets in trials with correct responses were assigned individual regressors and one regressor was assigned for all the cues and targets with incorrect responses. The regressors were modelled in the conventional GLM framework using custom MATLAB scripts developed within SPM toolbox. Single trial BOLD responses so estimated were used for the multi-voxel pattern analysis.

### MVPA analysis

MVPA analysis was performed by using linear support vector machine (SVM) implemented in the Statistics and Machine Learning toolbox of MATLAB. Trials formed the instances and the single-trial beta estimates of voxels within a given ROI as features. For feature attention, decoding was between attend-red vs attend-green; for spatial attention, decoding was between attend-left vs attend-right. A ten-fold cross validation technique was applied to determine the classification or decoding accuracy. The cross-validation analysis with different fold partition was repeated 25 times to avoid any intrinsic bias which may have resulted when dividing the trials into 10 specific groups (10-fold). Twenty-five, 10-fold cross validation accuracies (a total of 250 decoding accuracies) were averaged to obtain the classification accuracy for a subject. Group level accuracy was determined by averaging the classification accuracy across all subjects. Whether decoding accuracy in a ROI was above chance level was tested using a non-parametric Wilcoxon signed rank test. The resulting p values were adjusted for multiple comparisons by using the FDR technique (Benjamini and Hochberg 1995).

### Beta-series connectivity analysis

Single-trial level beta series was also used to perform functional connectivity analysis. The purpose of this analysis was to test whether cue-related functional connectivity between IFJ and V4 was modulated by the type of anticipatory attention (feature vs. space). Specifically, functional connectivity analysis was performed at the subject level by averaging the beta values over the voxels within a given ROI and subjecting them to a Pearson cross correlation analysis across trials. For attending feature, both attend-red and attend-green trials were combined; similarly, for attending space, attend-left and right trials were combined. The group level connectivity measure was computed by averaging the individual subject correlation coefficients after subjecting them to a Fisher’s (r to z) transformation.

## RESULTS

### Behavioral analysis

Behavioral data are shown graphically in Figures 2A and 2B, and the means and standard errors are provided in Table 1. A one-way ANOVA comparing the RT and accuracies (separately as factors) among the different trial types (attend left, attend right, attend red, and attend green) showed that no significant effects were observed between attention conditions for either RT or accuracy [F_RT_ (3,76) = 0.88, p = 0.45, Figure 2A; F_acc_ (3,76) = 0.72, p = 0.54, Figure 2B].The effect of attentional cueing on behavioral performance was tested by comparing the reaction time of validly and invalidly cued trials. Subjects responded more rapidly for validly cued trials than invalidly cued trials for both feature attention (t = 6.7, p<10^−6^; Figure 2C), and spatial attention (t = 4.8, p<10^−4^; Figure 2D). These results provide behavioral evidence that subjects deployed covert attention to the cued sensory attributes according to instructions and the level of attention was similar among different trial types.

**Figure 2.**
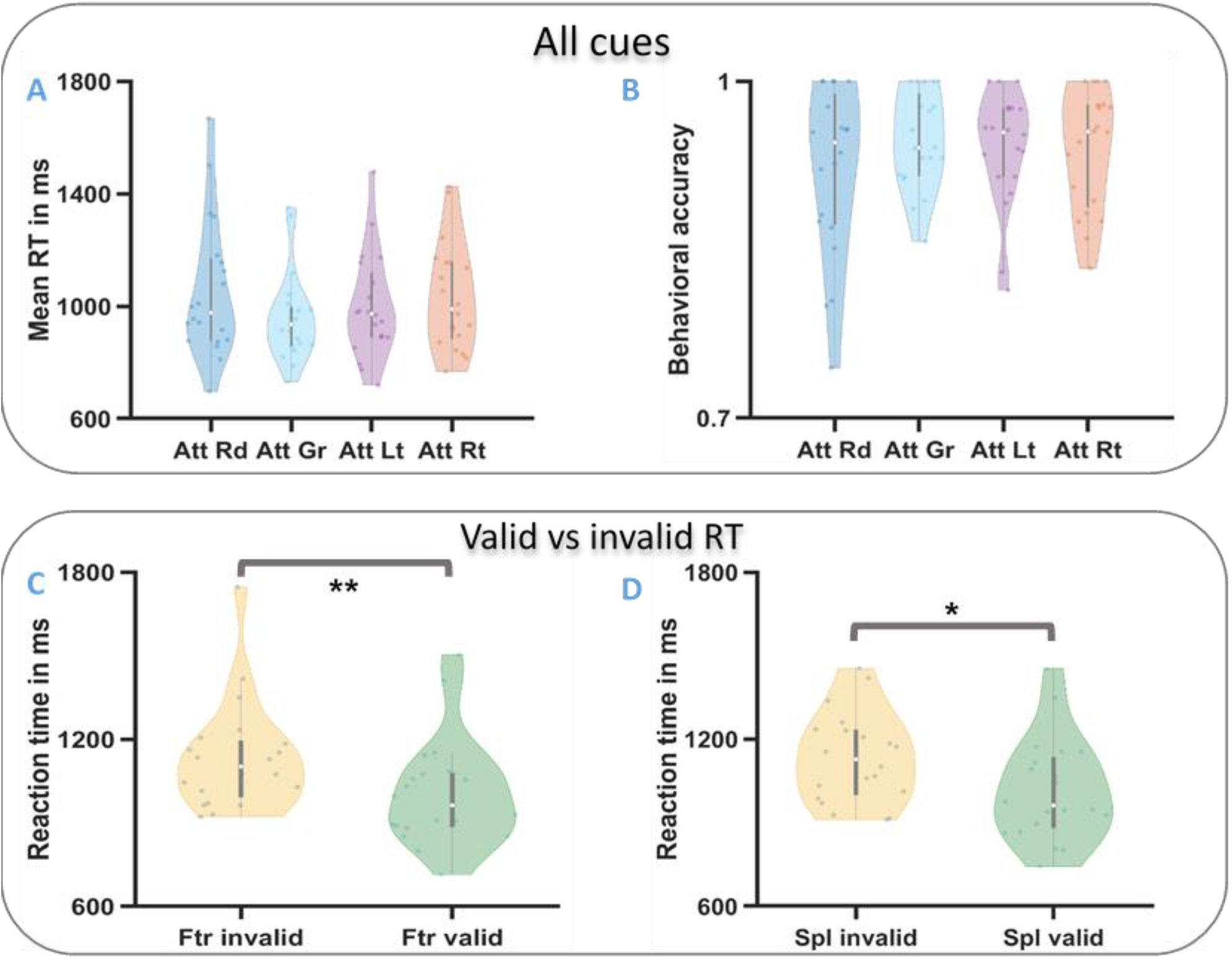
Distribution plots of behavioral performance during the task. (2A&2B): Pairwise comparison of reaction time (RT; 2A) and accuracies (percentage of correct trials; 2B) amongst different attention conditions. No significant behavioral differences were observed between the trial types. (2C &2D): Comparison of RTs for valid an invalidly cued trials, the differences were observed to be significant for both feature (2C; ** p<10^−5^) and spatial trials (2D; * p<10^−3^)).

**Table 1.** Reaction times (in ms) and accuracies (in %) for different attention conditions in the experiment. The values in the parenthesis indicate standard error of the mean.

| Behavior measure | Cue validity | Feature attention |  |  | Spatial attention |  |  |
| --- | --- | --- | --- | --- | --- | --- | --- |
|  |  | Attend red | Attend green | Mean | Attend left | Attend right | Mean |
| RT in ms | Valid | **1054**<br>(54.47) | **960**<br>(35.29) | **1006**<br>(44.80) | **1000**<br>(41.20) | **1032**<br>(42.31) | **1016**<br>(41.74) |
|  | Invalid | **1219**<br>(59.58) | **1303**<br>(62.75) | **1261**<br>(61.16) | **1016**<br>(40.84) | **1249**<br>(46.20) | **1132.5**<br>(43.52) |
| Accuracy in % | Valid | **92.20**<br>(1.6) | **94.60**<br>(1.0) | **93.40**<br>(1.3) | **94.00**<br>(1.2) | **93.90**<br>(1.2) | **93.95**<br>(1.2) |
|  | Invalid | **94.40**<br>(2.9) | **88.90**<br>(4.2) | **91.65**<br>(3.5) | **91.70**<br>(1.7) | **91.60**<br>(1.7) | **91.65**<br>(1.7) |

### Univariate analysis of cue-evoked BOLD activity

Univariate analysis of cue-evoked responses was analyzed using the GLM method. As shown in Figure 3, both color cues, spatial cues, and space + color cues activated the DAN (Figure 3A, 3B, and 3C), as well as bilateral IFJ (Figure 3D, 3E, and 3F), consistent with previous reports (Hopfinger et al., 2000; Corbetta et al., 2000; Giesbrecht et al., 2003; Slagter et al., 2007). Other regions activated included bilateral temporal cortex, bilateral dorsal anterior cingulate cortex (dACC), bilateral anterior insula, bilateral precuneus, bilateral inferior frontal gyrus (IFG), bilateral putamen, bilateral lingual gyrus, bilateral thalamus left supplementary motor area (SMA) (see Table 2). IFJ ROIs used in the subsequent analysis are shown in Figure 3G. A ROI-based univariate analysis revealed that IFJ activations were not significantly different between spatial cues and color cues (Figure 3H).

**Figure 3.**
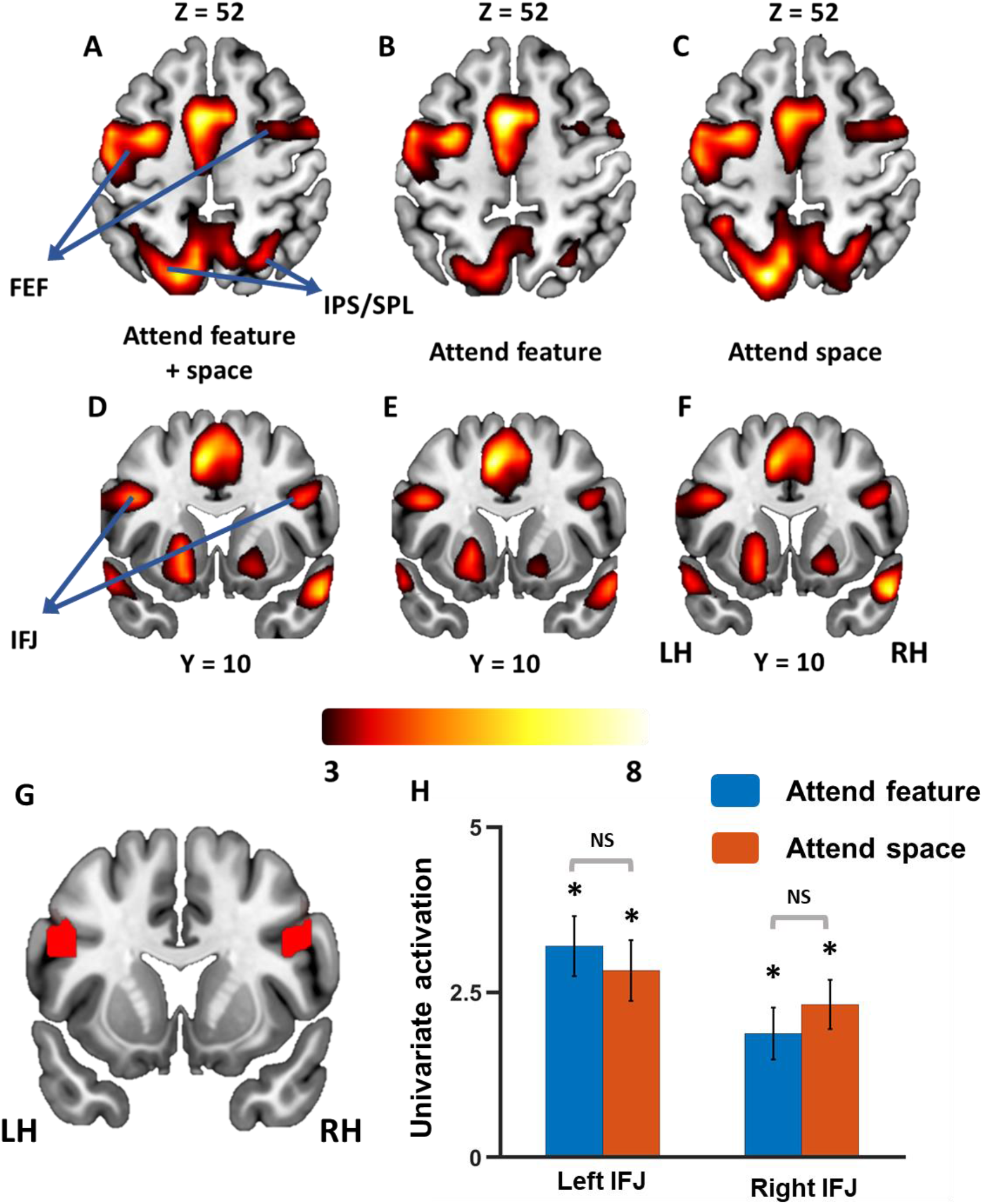
Univariate analysis of cue-evoked activity. (A, B & C): Both spatial cue and color cues activate dorsal attention network (FEF and IPS/SPL) (p<0.05, FDR corrected). (D, E & F): Both spatial cue and color cues activate bilateral IFJ (p<0.05, FDR corrected). (G): IFJ ROI defined based on activation by spatial + color cues. (H): Average univariate BOLD activation across subjects in left and right IFJ was compared between attending space and attending color. * indicates ROI activations above p < 0.05 (FDR). Error bars indicate SEM across subjects.

**Table 2.** MNI coordinates and corresponding Z scores are listed for brain areas showing significant BOLD activations for both spatial and feature cues. The group level activation maps were thresholded at p < 0.05 (FDR).

| Anatomical regions | Hemisphere | MNI coordinates<br>(x, y, z) | Z-score |
| --- | --- | --- | --- |
| Frontal eye field | Left | -27, -1, 52 | 4.8 |
|  | Right | 36, 2, 49 | 3.4 |
| IPS/SPL | Left | -15, -70, 52 | 4.9 |
|  | Right | 27, -61, 55 | 3.7 |
| IFJ | Left | -42, 11, 25 | 4.4 |
|  | Right | 42, 11, 25 | 4.2 |
| Temporal cortex | Left | 60, -19, -2 | 6.9 |
|  | Right | 63, -28, -2 | 6.5 |
| dACC | Left | -6, 5, 35 | 5.1 |
|  | Right | 6, 20, 37 | 3.5 |
| Anterior Insula | Left | -30, 26, 1 | 5.3 |
|  | Right | 33, 29, -4 | 4.7 |
| Putamen | Left | -18, 8, -5 | 4.5 |
|  | Right | 21, 14, -8 | 3.7 |
| Thalamus | Left | -9, -16, -2 | 4.8 |
|  | Right | 9, 16, 1 | 4.7 |
| Lingual | Left | -18, -79, 14 | 3.7 |
|  | Right | 12, -76, -5 | 3.0 |
| Inferior frontal gyrus | Left | -42, 11, 28 | 4.3 |
|  | Right | 45, 14, 25 | 4.2 |
| Precuneus | Left | -6, -64, 49 | 4.7 |
|  | Right | 3, -55, 46 | 4.0 |
| SMA | Left | -6, 5, 55 | 5.1 |

### Multivariate analysis of cue-evoked BOLD activity in IFJ

Cue-evoked beta series data from within right and left IFJ were subjected to MVPA analysis (See Figure 4). The decoding accuracies for attend red versus attend green were : 56.92% ± 1.5% (right IFJ), and 57.41% ± 1.5% (left IFJ), which were significantly above the chance level of 50% (p<0.05 FDR corrected) (Figure 4A). The decoding accuracy for attend left versus attend right, however, were not significantly above chance level (rIFJ: 52.96 ± 1.7 %, p=0.17 (FDR); lIFJ: 52.00 ± 1.8 %, p=0.39 (FDR) (Figure 4B). It is evident that multivoxel patterns clearly distinguished different attended information for feature attention but not for spatial attention, even though both forms of attention activated the IFJ ROI in the univariate analyses (Figure 3). These results support the idea that IFJ is more involved in the control of feature attention than spatial attention.

**Figure 4.**
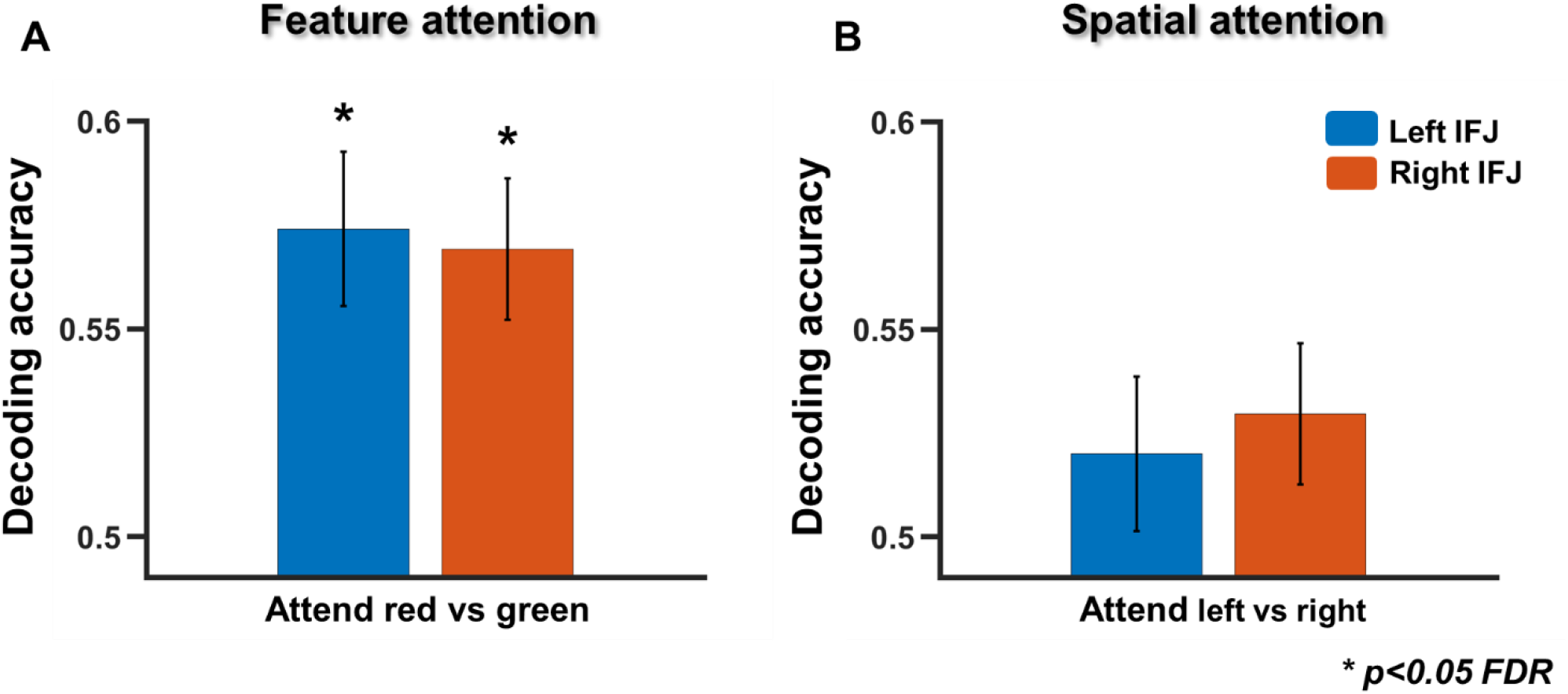
Multivariate analysis of cue-evoked BOLD activation in IFJ. Decoding accuracies between attend red vs attend green (Feature; Figure 4A) and between attend left vs attend right (Spatial; Figure 4B) in left and in right IFJ. Decoding accuracy for attend red versus attend green was significantly above chance level in both left sand right IFJ, whereas decoding accuracy for attend left versus attend right was not significantly above chance level in either left or right IFJ. *p<0.05 FDR. Error bars indicate SEM across subjects.

### Analysis of target-evoked activity in IFJ

To what extent is IFJ involved in attentional selection of the target stimulus in visual cortex? To address this question, we analyzed the target-evoked beta series data. Both univariate activation and MVPA decoding analysis were performed for attend red versus attend green, and for attend left versus attend right. As shown in Figure 5A, both IFJs were activated by the target stimuli (p<0.05 FDR) irrespective of whether the attention was paid to space or color, and the activations are not different between attend space and attend color conditions. Similarly, MVPA decoding analysis showed that the decoding accuracies for target-evoked activity in the IFJ were not above chance level for feature attention (attend red vs. attend green) (rIFJ: 54.6% ± 2%, p = 0.1; lIFJ: 53.4 % ± 1.8%, p = 0.1) or for spatial attention (attend left vs. attend right) (rIFJ: 51.6% ± 2%, p = 0.6; lIFJ: 52.20 % ± 1.7%, p = 0.6). Effect sizes comparing the decoding accuracies from cue and target-evoked data were observed to be small for feature attention (rIFJ: d = 0.24; lIFJ: d = 0.47), whereas for spatial attention there were no effects (rIFJ: d = 0.12; lIFJ: d = - 0.02). These results suggest that IFJ is not strongly involved in attentional selection of targets for feature attention.

**Figure 5.**
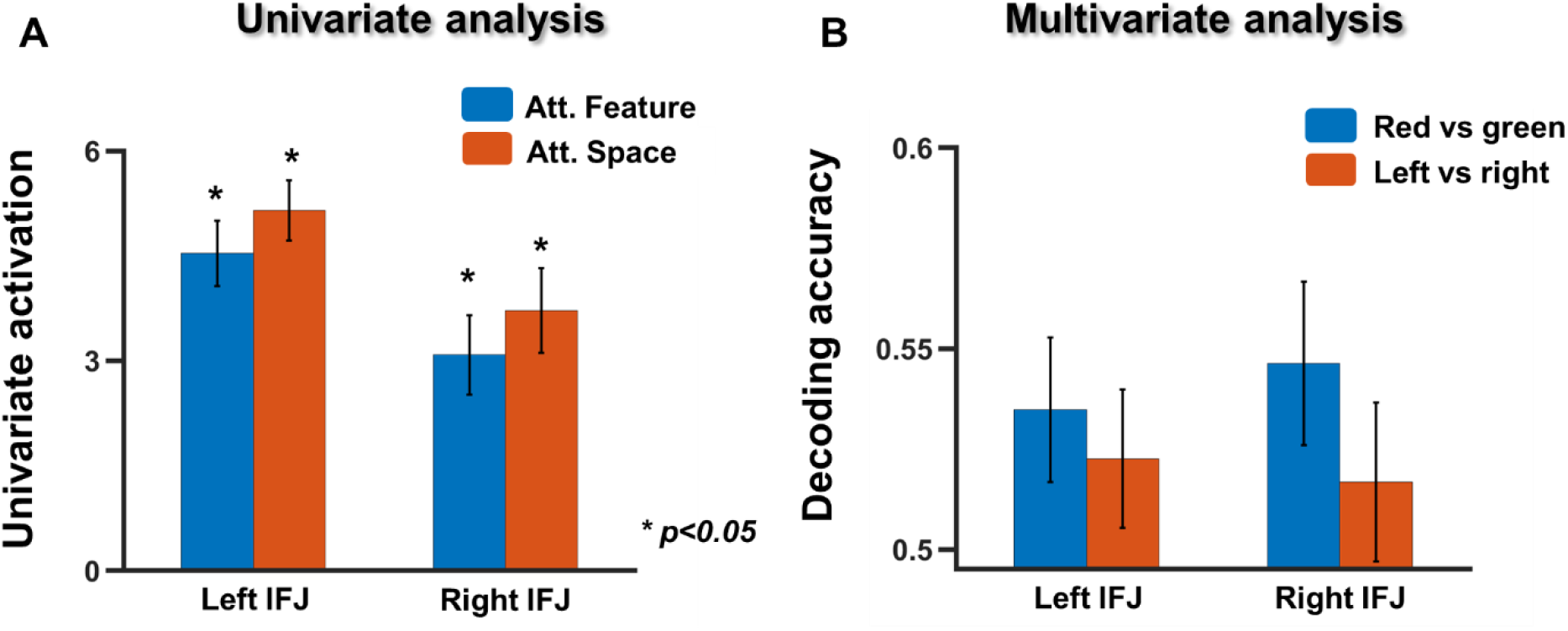
Target-evoked activity in the IFJ. Univariate and multivariate analysis of target-evoked activity in IFJ. (A): Univariate activation. (B): MVPA decoding accuracies for attend red vs green (feature) and attend left vs right (spatial); neither were significantly above chance decoding. Error bars indicate SEM across subjects.

### Connectivity between IFJ and V4 in attentional control and selection

During covert attention, biasing signals issued by attention control regions such as IFJ propagate to the sensory regions to influence the processing of the upcoming stimulus (Desimone and Duncan 1995; Geisbrecht et al., 2003; Giesbrecht et al., 2006; Slagter et al., 2007). A visual sensory area that has been extensively investigated for its role in processing color information is V4 (Lueck et al., 1989; Zeki et al., 1991; Murphey et al., 2008; Zanto et al., 2010). How does IFJ interact with V4 during the cue-to-target interval? What are the behavioral consequences of such interaction? We investigated these issues by treating bilateral V4 as a single ROI and computing the functional connectivity between V4 and right IFJ, and separately between V4 and left IFJ (Figure 6A). Left IFJ did not show significant functional connectivity for either color cues (p<0.16) or spatial cues (p<0.8) (Figure 6B). In contrast, right IFJ (rIFJ) connectivity with V4, following color cues was found to be significantly greater than zero (p<0.00001), whereas the functional connectivity between rIFJ and V4 for spatial cues was not significantly different from zero (p<0.23). The functional connectivity for rIFJ-V4 following color cues was also significantly greater than that following spatial cues (p<0.05) (Figure 6C). Behaviorally, for subjects with higher performance accuracy, rIFJ-V4 functional connectivity was higher; see Figure 6D.

**Figure 6.**
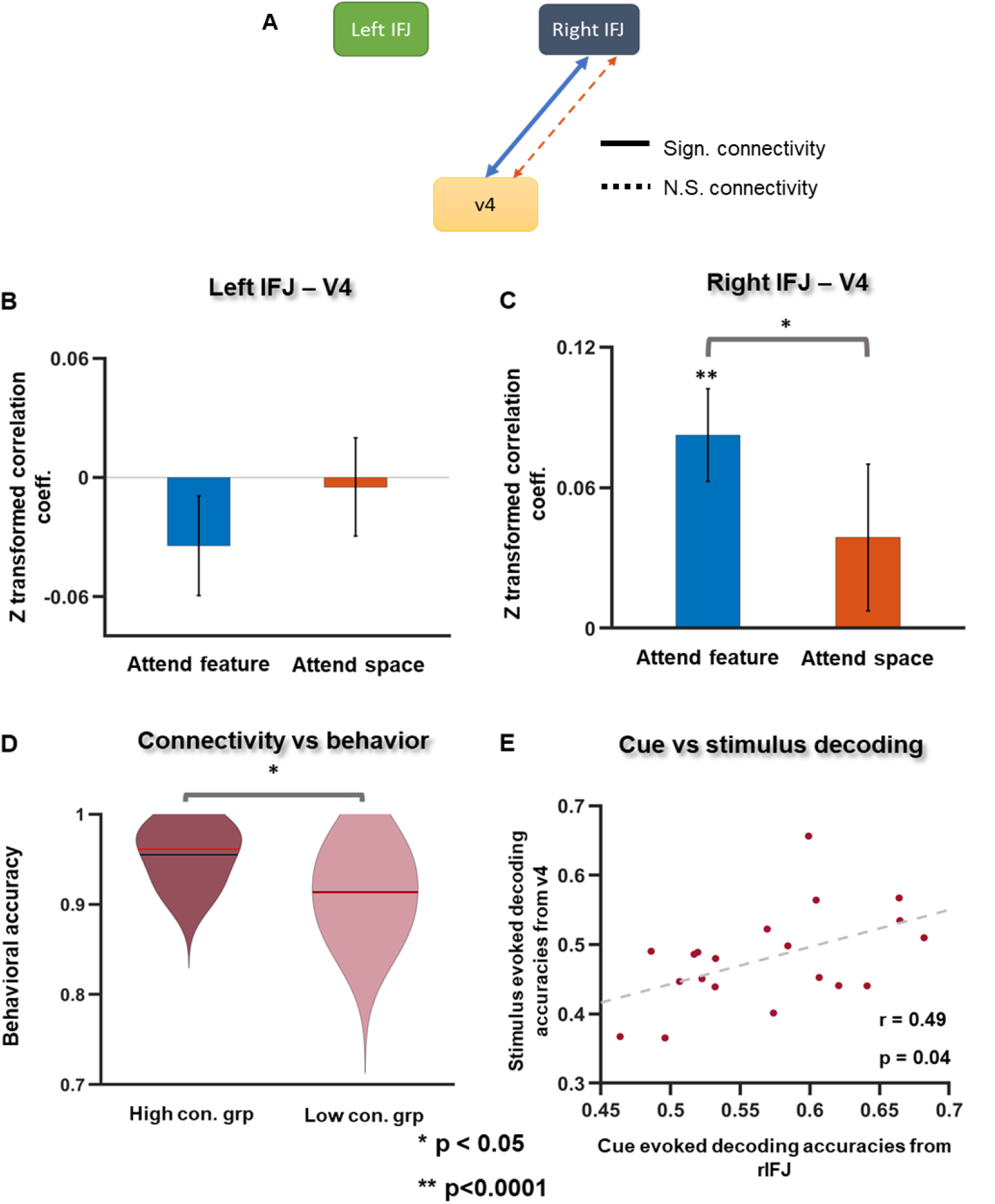
Relationship between IFJ and V4. (A): Schematic representation of connectivity between IFJ and V4. Here the solid lines denote significant functional connectivity and the dashed lines represent non-significant functional connectivity between IFJ and V4; the thickness of the solid lines denotes its strength. (B&C): Cue-related functional connectivity between IFJ and V4 for attend feature and attend space trials. IFJ-V4 functional connectivity during feature attention was significantly higher compared to spatial trials for right IFJ (right), while left IFJ showed no significant functional connectivity for either types of trials. (D): Subjects with higher behavioral accuracies had stronger rIFJ-V4 functional connectivity. (E): Correlation between cue-evoked decoding accuracies in rIFJ and target-evoked decoding accuracy in V4.

The prevailing model is that preparatory attention signals in IFJ influence target processing of visual cortical areas such as V4. To test whether IFJ enhances attentional selection of targets in V4 we applied MVPA to the target evoked beta series data within V4. The decoding accuracy between selecting the red target versus selecting the green target in V4 was correlated with the decoding accuracy of attend red versus attend green in rIFJ (r = 0.49, p = 0.04; Figure 6E). A similar comparison for left IFJ did not result in significant correlations (r =0.32 p = 0.18), suggesting that the strength of the attention control signals in rIFJ, rather than lIFJ, predicted the strength of attentional selection of the stimulus in V4.

## DISCUSSION

The role of IFJ in attentional control was examined in this study. Four questions were considered: (1) Is IFJ’s role in attention control specific to feature attention? (2) Is there a hemispheric difference in IFJ control? (3) Is IFJ involved in both attention control and attention selection? (4) What is the relation between IFJ and sensory area V4 in feature (color) attention? We recorded fMRI data from subjects performing a cued spatial and non-spatial (feature) attention experiment, in which during spatial attention trials they were cued to attend to a location in either the right or left visual field independent of stimulus color, whereas in feature attention trials they were cued to either attend to red or green stimuli, independent of stimulus location.

Conventional univariate fMRI analysis found that both spatial cues and color cues evoked significant preparatory attention-related BOLD activity in bilateral IFJ, as has been observed previously (Giesbrecht et al., 2003). Moreover, attend left and attend right cues evoked similar univariate BOLD activation in IFJ as attend red and attend green cues. Multivariate pattern analysis was then applied, and decoding accuracy for feature attention (cue red vs. cue green) in both left and right IFJ was significantly above chance level. However, for spatial attention (cue left vs. cue right) decoding was at chance level in both left and right IFJ, suggesting that cue related IFJ activity is more specifically geared toward feature attention control rather than spatial attention control. MVPA analysis of target evoked data further showed that neural patterns within IFJ did not distinguish between attentional selection of the left target vs the right target. Decoding accuracy of attentional selection between the red target vs the green target was slightly higher, but it also did not reach statistical significance, although it had attained a small effect size. These results suggested that IFJ is mainly involved in attentional control rather than attentional selection. During feature cue processing, the right IFJ showed enhanced functional connectivity with the sensory area V4, and this enhanced connectivity was positively associated with behavior; the higher the rIFJ-V4 functional connectivity, the better the behavioral performance. Moreover, the decoding accuracy between cue red versus cue green (cue processing) in IFJ predicted the decoding accuracy between selecting red versus selecting green (target selection) in V4.

### IFJ as a region for attention control

Giesbrecht et al., (2003) and Slagter et al., (2007) were among the first to report activation of regions in and around IFJ along with the DAN during both spatial and feature attentional control (using visually presented cues). Based on the resting state connectivity Corbetta et al. (2008) later considered IFJ (labeled as MFG in their report) part of the attention control networks of the DAN and VAN (He et al., 2007). They hypothesized that IFJ may act as a relay station which sends the top-down signals from DAN to visual cortex during endogenous attention and re-orienting signals towards salient stimuli from VAN to DAN during exogenous attention. A study by Tamber-Rosenau et al. (2018) investigated whether IFJ plays a role in maintaining sustained attention along with DAN during spatial attention. Participants, following a color cue, directed their attention to one of the four peripheral visual fields and anticipated the processing of an impending target. Analyzing event-related BOLD activity, they reported significant activity in bilateral IFJ ROIs immediately after the cue as well as after target onset, but no significant delay activity in the cue-to-target interval, a pattern which is in contrast to that observed in the IPS and FEF of the DAN, which did show significant sustained BOLD activity in the cue-to-target interval. They thus concluded that IFJ activity is transient, extending earlier work by Asplund et al., 2010, and is involved in interpreting the cue and orienting attention towards the attended hemifield, rather than in controlling covert attention, with the latter being carried out mainly by the DAN. In our data, spatial cues elicited significant activity within the IFJ, but attend left and attend right cues are not distinguished in either the univariate (results not shown) or the multivariate analysis. An analysis of the same data revealed significant MVPA decoding of attend left versus attend right cues in the DAN (Supplementary Fig. 1 from Rajan et al., under review at Journal of Cognitive Neuroscience). These results are consistent with the interpretation that IFJ is engaged transiently following the spatial cues but not involved in maintaining covert spatial attention in the cue-to-target interval; the latter was carried out by DAN instead.

### Role of IFJ in controlling top-down attention to feature

Zanto et al. (2010) suggested that IFJ is involved in the control of feature attention. They asked the participants to view and remember 4 different sets of dots during encoding (2 each for different colors and motion), and to report during retrieval whether the target was present in one of the two stimuli encoded in the attended feature domains (color or motion). Using functional connectivity as the observable, and visual cortex ROIs as seed regions (V4 for color and MT for motion), they showed that during encoding, IFJ voxels exhibit stronger IFJ-V4 connectivity when attending color, and stronger IFJ-MT connectivity when attending motion. Baldauf and Desimone (2014) presented face-house compound stimuli to subjects in a one-back paradigm and asked the subjects to either attend faces or attend houses. Using gamma frequency synchrony as the measure, they found increased functional interaction between IFJ and FFA when attending faces and increased functional interaction between IFJ and PPA when attending houses, thus extending the work of Zanto and colleagues to object attention. We point out that in Zanto et al. (2010) and Baldauf and Desimone (2014), attention was directed to different feature domains but not to subcategories of stimuli within a feature domain. It is therefore not clear whether IFJ activity codes different attended information in a given feature domain.

Recent work by Gong and Liu (2020) asked participants to pay attention to one of two subcategories of stimuli within a feature domain: e.g., clockwise versus anti-clockwise motion in the motion domain, or red versus green in the color domain. The participants’ task was to detect brief changes in luminance in colored moving dots in the attended feature domain. Multivoxel patterns evoked by attention to different stimuli within a given feature domain were found to be decodable in IFJ. While our results showing that IFJ decoded attended color information in the cue-to-target interval are consistent with Gong and Liu (2020), our study differs from theirs in two important respects. First, in their study, a stimulus (colored moving dots) was present throughout the trial, and the subject were to detect a change in the stimulus. As a result, there was no distinction made between preparatory attentional control and selective target processing. By having a cue-to-target interval in our paradigm it was possible to isolate attentional control from stimulus selection, and therefore conclude that the IFJ is involved in controlling attention to stimulus features (in the absence of sensory input) but not in attentional selection of target stimuli (see below). Second, by including both attention to feature and attention to space in one paradigm, we are also able to conclude that IFJ plays a more significant role in the control of feature attention than the control of spatial attention.

### Hemispherical differences in IFJ control of attention

Many brain functions are lateralized in humans (Gazzaniga, 2015). Prominent examples include language (left lateralized) and spatial cognition (right lateralized). Is there a hemispheric asymmetry in IFJ control of feature attention (Asplund et al., 2010; Zanto et al., 2010, Zanto et al., 2011, Baldauf and Desimone 2014, Tamber-Rosenau et al., 2018, Zhang et al., 2019)? Zanto et al. (2011), using rTMS, showed that stimulation of the right IFJ impaired attention to color where stimulation of both left and right IFJ impaired attention to motion, suggesting a right lateralization of attention to color but not attention to motion. In contrast, other studies have found no differences in functionality between the left versus right IFJs. Zhang et al. (2018) directed subject’s attention to either motion or color in laterally presented visual stimuli and reported increased activity within bilateral IFJ. When the authors divided the hemispheres into contralateral and ipsilateral based on the spatial location of the stimuli, they found the contra-lateral IFJ to be more involved in directing the global effect of the feature attention. Bilateral IFJ activity in terms of MVPA decoding was also observed in (Gong and Liu 2020). In our data, univariate activation and MVPA decoding results are consistent with previous reports, namely, in both IFJs, there are similar univariate activations for red and green cues, and attend red and attend green are equally well decoded in the left and the right IFJ. However, hemispheric differences emerged with the application of techniques other than univariate activation and MVPA decoding. Applying functional connectivity analysis, we found that right IFJ-V4 connectivity is significant in feature trials and higher than that in spatial trials. This enhanced rIFJ-V4 functional connectivity predicted better behavioral performance; left IFJ-V4 connectivity is not significant in both types of trials. This result is in line with the findings of Zanto et al. (2010), indicating that attention to color is more right lateralized in IFJ when IFJ’s interaction with sensory cortex is taken into account (Sasaki et al., 2007; Ikegami et al., 2012; Takano et al., 2014). Furthermore, the cue-related decoding accuracy in right IFJ, not in left IFJ, predicted the efficacy of target selection in V4 (see Figure 6). Taken together, these findings suggest that while both IFJ contained attention control signals, it is the right IFJ that plays a more prominent role in influencing sensory brain regions during covert feature attention.

### The role of IFJ in attentional selection

Both the extant literature and the findings reported here support the notion that IFJ is involved in the preparatory control of feature attention. Whether IFJ plays a role in attentional selection of the target stimuli remains less clear. Baldauf and Desimone (2014) observed an increase in BOLD activity around right IFJ when participants-attended to the color (red or green) of moving dots compared with passive viewing of moving dots. Bichot et al. (2015) observe an increase in firing rates in neurons in VPA (the non-human primate analog of IFJ) when the attended stimulus was the primary target in a visual search task. IFJ is sometimes considered to be at the confluence of dorsal and ventral attention networks, dynamically switching its association between DAN and VAN during anticipatory attention versus stimulus selection processes, as has been shown by He et al., 2007 (who labeled it MFG), and others (Apslund et al., 2010; Tamber Rosenau et al., 2018). Whereas IFJ activation by target stimuli was expected, whether it is engaged in selecting the attended stimulus over the unattended stimulus within the same feature domain remains to be tested. In our data, univariate analysis revealed activation by red and green targets in bilateral IFJ. Decoding selecting red versus selecting green was also found not to be significantly above chance, thus indicating that IFJ is more involved in exerting top-down control and sensory modulation following cue processing and in anticipation of the upcoming target, rather than in active selection of the target stimuli.

### Summary

In this study, using univariate and multivariate analysis, we provide evidence to support the idea that IFJ is involved in the control of feature attention rather than spatial attention. The right IFJ plays a more significant role when the relation between IFJ and sensory areas are considered using functional connectivity analyses. Moreover, it appears that IFJ mainly functions as a controller of covert attention rather than the selector of attended stimuli.

## Acknowledgements

This work was supported by NIMH grant MH117991 (G.R.M. and M.D.). All data will be publicly available on the NIMH Data Archive.

