## Supplementary Figure 1 for "Role of Inferior Frontal Junction (IFJ) in the Control of Feature vs Spatial Attention"

### Top-down control of spatial and feature attention within DAN

To test if voxel patterns within DAN predict the attended condition, we performed MVPA analysis for attend red versus green as well as for attend left versus right by considering voxels within DAN (Figure S1A). The classifier performance was derived and tested for chance level at $p<0.05$ (Wilcoxon signed rank test). We found significantly above chance level decoding for both feature (attend red vs green) and space (attend left vs right) (Figure S1B). Further, decoding accuracies between attend red vs attend green in DAN and in IFJ are significantly correlated (Pearson correlation $r = 0.78; p=10^{-6}$), indicating that these regions may work in tandem to effect top down control of feature- based attention (Figure S1C).

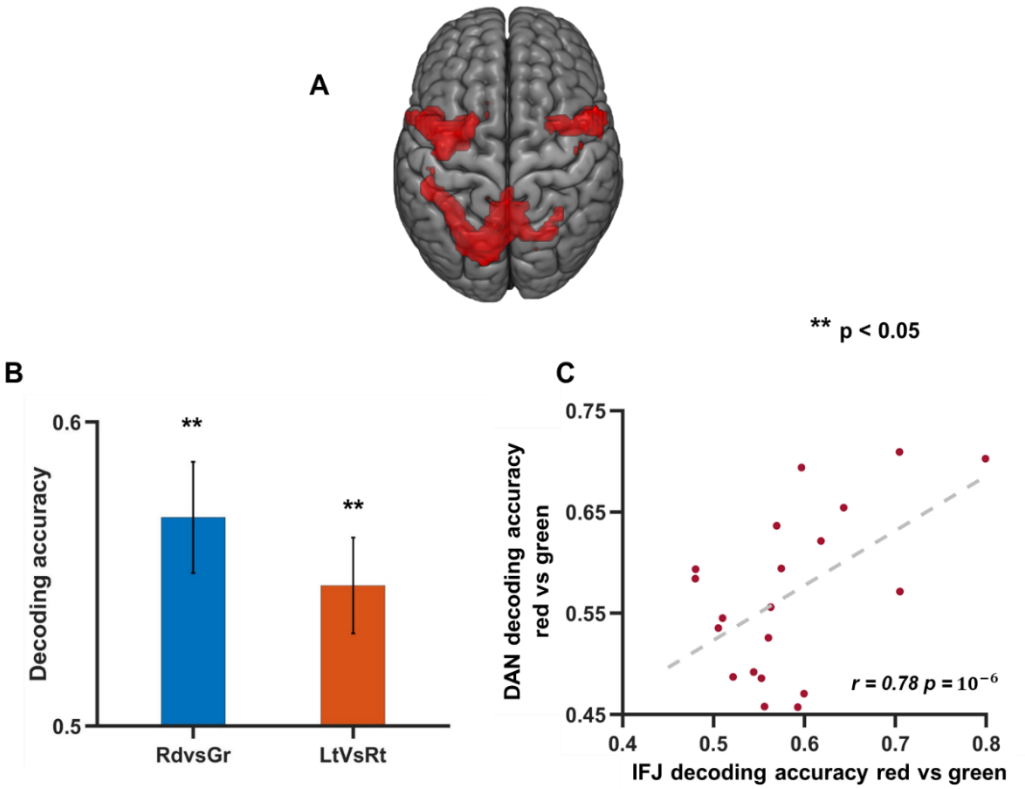


**Figure S1**. Decoding accuracy comparison between feature attention (attend red vs attend green) and spatial attention (attend left vs attend right) in the DAN using cue evoked neural patterns. **A**: The DAN ROI. **B**: Decoding accuracies for feature and spatial attention cues. **C**: scatter plot comparing decoding accuracy within bilateral IFJ and DAN for attend red vs green.
